# Intraspecific bacterial competition mediated by rapidly diversifying tailocin and prophage loci

**DOI:** 10.1101/2025.04.19.648921

**Authors:** Sarah J. Kauffman, Ryan M. Awori, Emmanuel C. Allwell, Aaron Taylor, Farrah Bashey, Heidi Goodrich-Blair

**Author notes:** Correspondent footnote. These authors contributed equally to the work (Sarah J. Kauffman, Ryan M. Awori).

## Abstract

Signatures of selection in microbial genomes are often linked to biotic interactions, notably resistance to host immunity or bacteriophage attack. Here, we highlight the importance of competitive interactions, specifically between conspecific bacteria, in shaping microbial genomes, using animal-associated *Xenorhabdus* bacteria. An aspect of microbial genome variation is the presence of diverse mobile genetic elements that distinguish bacterial genomes from their closely related kin. We found that compared to those across domain *Bacteria*, *Xenorhabdus* genomes contain among the highest proportion of phage-related genes, and that variation among strains in their total number of protein-coding genes is largely predicted by variation in total number of non-cargo phage genes per genome. A universal yet highly variable *Xenorhabdus* phage-related region encodes xenorhabdicin tailocins. This region ranged in length from 12 to 41 kilobases, and its specificity-determining main tail fiber varied from 341 to 1035 amino acids. Concomitant with this variation, tailocins produced by six strains of *X. nematophila* differed dramatically in particle length and killing profile. Intriguingly, while *X. nematophila* xenorhabdicins displayed common heterospecific killing activity, they varied in conspecific killing activity. We further demonstrate the ecological importance of xenorhabdicin diversity by associating intraspecific variation in killing profiles of mitomycin-induced lysates from 42 sympatric *X. bovienii* strains with genes from both xenorhabdicin-encoding loci and prophages. The susceptibility profiles of strains to lysates were associated with O-antigen biosynthesis genes. Overall, our data demonstrate that through bacteriophage-mediated genome diversification, an animal-associated bacterium can tailor its weaponry and defense systems to target the closest of relatives.

**IMPORTANCE:** Microbes exist in complex communities with each other and with animal or plant hosts. Such biotic interactions can impose strong and variable selection on microbes and are predicted to foster diversity within lineages. By examining genomes from one bacterial genus, we show that phage-associated genes are responsible for an unusually high degree of variation in protein-coding gene content. Moreover, we link this variation to functional diversity in competitive interactions among *Xenorhabdus* strains. Different strains of *Xenorhabdus* bacteria frequently co-infect an insect, which simultaneously promotes strong selection for competitive dominance within an insect host as well as opportunities for horizontal gene flow. Additionally, genomic rearrangement and homologous recombination can provide phenotypic variation for selection. The tractability of *Xenorhabdus* for both lab and environmental studies make them powerful for understanding microbial genome evolution and competition and this study reveals the dominant role of bacteriophage and bacteriophage-like elements in these processes.

## INTRODUCTION

Diversification and adaptation of microbial lineages is facilitated by their flexible genomes. Variation arises through horizontal gene transfer as well as rearrangements induced by insertion sequences and transposons (1, 2). Hot spots of diversification are often linked to bacterial interactions either with their eukaryotic hosts or bacteriophage predators (3–5). Less appreciated as a force of diversification are the interactions among microbes themselves (6). Microbes have myriad machineries that are deployed during competition with each other (7). These range from small diffusing molecules to elaborate phage-derived nanoparticles. A single bacterium may have an arsenal of attacking tactics thought to be important in different ecological contexts (8). Additionally, resistance to competitive weapons and the costs of resistance have been implicated in maintaining diversity within microbial populations (9–12). Understanding the impact of competitive interactions on the evolution of bacterial genomes within a taxon can not only provide an important source of novel anti-infectives but also afford insights into how to robustly manipulate microbial communities (13, 14).

*Xenorhabdus* bacteria are excellent models to understand the evolution of genomes within complex communities. *Xenorhabdus* are entomopathogenic bacteria that are mutualistically associated with *Steinernema* nematodes. In this obligate association, the non-feeding, infective juvenile stage of the nematode carries its specific *Xenorhabdus* bacterial symbiont into the insect prey that will be consumed. A primary service provided by *Xenorhabdus* is the killing and consumption of insect biomass, which they use to increase their own biomass, which is then consumed by the bacterivorous nematodes (15, 16). A second service provided by *Xenorhabdus* symbionts is the protection of the insect cadaver from competitors and predators, including fungi, ants, and other bacteria (17–22).

The *Xenorhabdus* life cycle can be roughly divided into three stages: a virulent stage, in which a small population of bacteria are released from the infective juvenile nematode into the insect blood cavity, where they suppress immunity and contribute to insect death; a growth phase in which the bacteria consume the insect tissues, reproduce and colonize the cadaver, while also being consumed by the developing nematodes; and the transmission stage, in which the bacteria colonize the intestine of the developing infective juvenile stage of the nematode (23, 24). This symbiotic lifestyle of repeated cycles of virulence, rapid growth, colonization, and persistence influence genome evolution (24). Experimentally, *X. nematophila* evolution within the context of mutualistic colonization of the transmission stage nematode constrains evolution of insect virulence phenotypes (25). The transmission stage also represents an obligatory and recurring potential bottleneck in *Xenorhabdus* genome diversity. Each *Steinernema* nematode species recruits, within its anterior intestine (26), a *Xenorhabdus* species for transmission to the exclusion of other *Xenorhabdus* species (27–29). Further, for the two species that have been tested, *X. nematophila (*ATCC19061 and AN6/1) and *X. bovienii* (SS-2004), the transmitted bacterial population derives from just 1-2 individual cells that grow to fill the receptacle, though this may not be universally true for all *Xenorhabdus* (28, 29). Finally, genome evolution is likely impacted by the obligate stage in which a small *Xenorhabdus* population persists in the absence of exogenous nutrients for up to many months within their infective juvenile nematode host before resuming their lifecycle within a new insect prey. The insect environment offers further selective pressures. Apart from *Steinernema hermaphroditum*, all other *Steinernema* studied to date require infection by both male and female nematodes. Since each individual nematode carries its own largely clonal *Xenorhabdus* population (30), every newly infected insect comes with the potential for competition between conspecific *Xenorhabdus* strains. The outcome of such within-host competition can have impacts on the fitness of the nematode-bacterium complex. For instance, *Xenorhabdus* strains which grow faster than or inhibit the growth of other strains *in vitro* are also competitively dominant *in vivo* to the benefit of their nematode host (31, 32). This competitive inhibition of conspecific strains comes through xenorhabdicin tailocins (20, 31, 32) whose corresponding genome loci contain highly variable regions, partly due to numerous receptor binding domain (RBD) encoding tail fiber genes (21).

In the current study, we sought to identify genomic aspects that predispose *Xenorhabdus* to flexibility and diversification within its complex niche. Further, we sought to link this genomic variation to the diverse, antagonistic competitive interactions we observed between conspecific strains.

## RESULTS

### Six percent of *Xenorhabdus* genomes is mobilome

To characterize diversification among *Xenorhabdus* genomes, we used the genomes of 97 strains from 27 species obtained from *Steinernema* isolated from around the world to create a pangenome that contained 381,400 protein-coding genes (Fig. 1, Supplementary File 1). The average genome length, guanine-cytosine (G+C) content, and the total number of protein-coding genes for a *Xenorhabdus* genome were 4.54 megabases (Mb), 44.33%, and 3,944, respectively. Notably, the ranges for the total number of protein-coding genes and genome length were 2,734 – 4,803 and 3.18Mb - 5.35Mb, respectively, suggesting considerable genome diversification within the genus.

**Figure 1.**
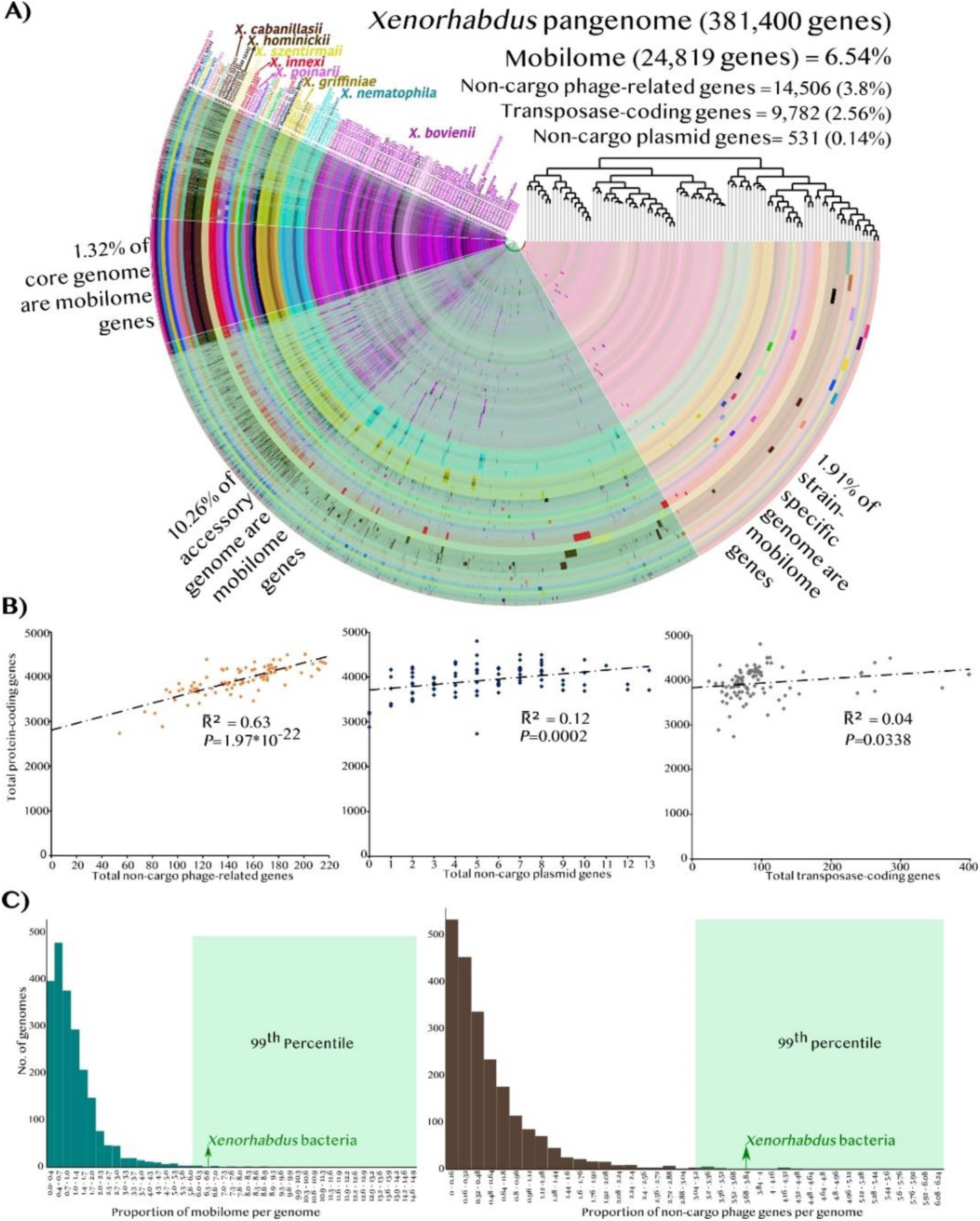
*Xenorhabdus* bacteria have comparatively large mobilomes, which are associated with genome diversification. A) Graphical representation of a pangenome of 97 strains of 27 species from the genus *Xenorhabdus*, and its mobilome proportion and distribution. Each of the 97 concentric arcs represents a genome. Each radius represents a group of orthologous genes (gene cluster). Dark shades along a radius indicate the presence of a gene cluster (GC) in the genome. The eight species with more than one strain in the pangenome are shown in large font. The pangenome was composed of 381,400 protein coding genes grouped into 15,644 GCs. There were 1,520 core, 9,562 accessory and 4,562 strain-specific GCs, where we defined core, accessory, and strain-specific GCs as those constituted of orthologs from n=97, n=2-96, and n=1 genomes, respectively. The dendrogram on the top left is based on hierarchical clustering based on presence and absence of GCs. B) Scatterplots depict correlations between mobilome and proteome (total number of protein-coding genes) sizes in 97 *Xenorhabdus* genomes. Each point in the graphs represents a genome. Sixty-three percent of the variation in total protein-coding genes in genomes was accounted for by the number of phage-related genes per genome. C) *Xenorhabdus* genomes have among the highest mobilome proportions among bacterial species whose genome sequences are known. The histograms depict the mobilome and non-cargo phage genes proportions in 2,196 circularized bacterial genomes of the NCBI COG 2024 database.

Characterization of the mobilome (Fig. 1A), revealed that *Xenorhabdus* bacteria were in the top 1% of strains whose genomes contain the largest mobilome proportions in general and specifically non-cargo phage genes, when compared to those of 2,196 similarly annotated complete bacterial genomes from the NCBI COG 2024 database (https://www.ncbi.nlm.nih.gov/research/cog-project/) (Fig. 1C). The coefficients of determination for a genome’s mobilome size to its length and proteome size, were 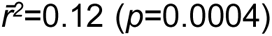 and 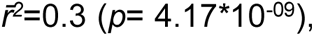 respectively. Mobilome category correlations (Fig. 1B), revealed that total number of phage-related genes in a *Xenorhabdus* genome accounted for 63% of the variation in the proteome size. This demonstrated a strong association between phage-related genes and genome diversification in the *Xenorhabdus* genus.

### The xenorhabdicin locus exemplifies phage-associated genome diversification

To find a few of the genomic loci of these phage-related genes, we identified eighty prophage/ phage-like regions (Supplementary File 1) in the ten most phylogenetically diverse, least-fragmented genome assemblies from our dataset. On average, each genome had eight prophages that accounted for 6.5% of the genome’s base pairs and contained 9.8% of its total protein-coding genes. The average number of prophages was in between previous estimates of six (33) and nine (34). Every genome also contained a phage-like locus that encoded xenorhabdicin tailocins.

Xenorhabdicin tailocins are extracellular rigid contractile phage tail-like structures with no toxin delivery mechanism but terminally puncture membranes of target bacteria (19–21, 35–38). Ecologically, xenorhabdicins kill co-infecting *Xenorhabdus* strains within the dead insect niche (31) to the advantage of the nematode host of the xenorhabdicin-producing population (20, 32, 39). Xenorhabdicin biosynthesis is encoded by *xnp1/xbp1* loci (19, 20) and for the 62 loci we analyzed, it ranged in length from 12.2 kilobases (Kb) in *X. japonica* to 41.3 Kb in *X. bovienii* LD8. This length variation was primarily due to differences in gene content in a highly variable region that lies between the *N*-terminal encoding main tail fiber gene fragment and the tail sheath gene (Fig. 2). Conversely, highly conserved genes included those encoding a spanin; transcriptional regulators AraC, OgrK, CI phage repressor; the SOS response regulator DinI; baseplate hub proteins BH1a, BH1 and BH2 (40); the baseplate wedge proteins BW1, BW2 and BW3 (40); and the tail spike (Fig. 2). Also present across all loci were the tail tube, tail sheath and tapemeasure genes, the last of which had a clear variation in length.

**Figure 2.**
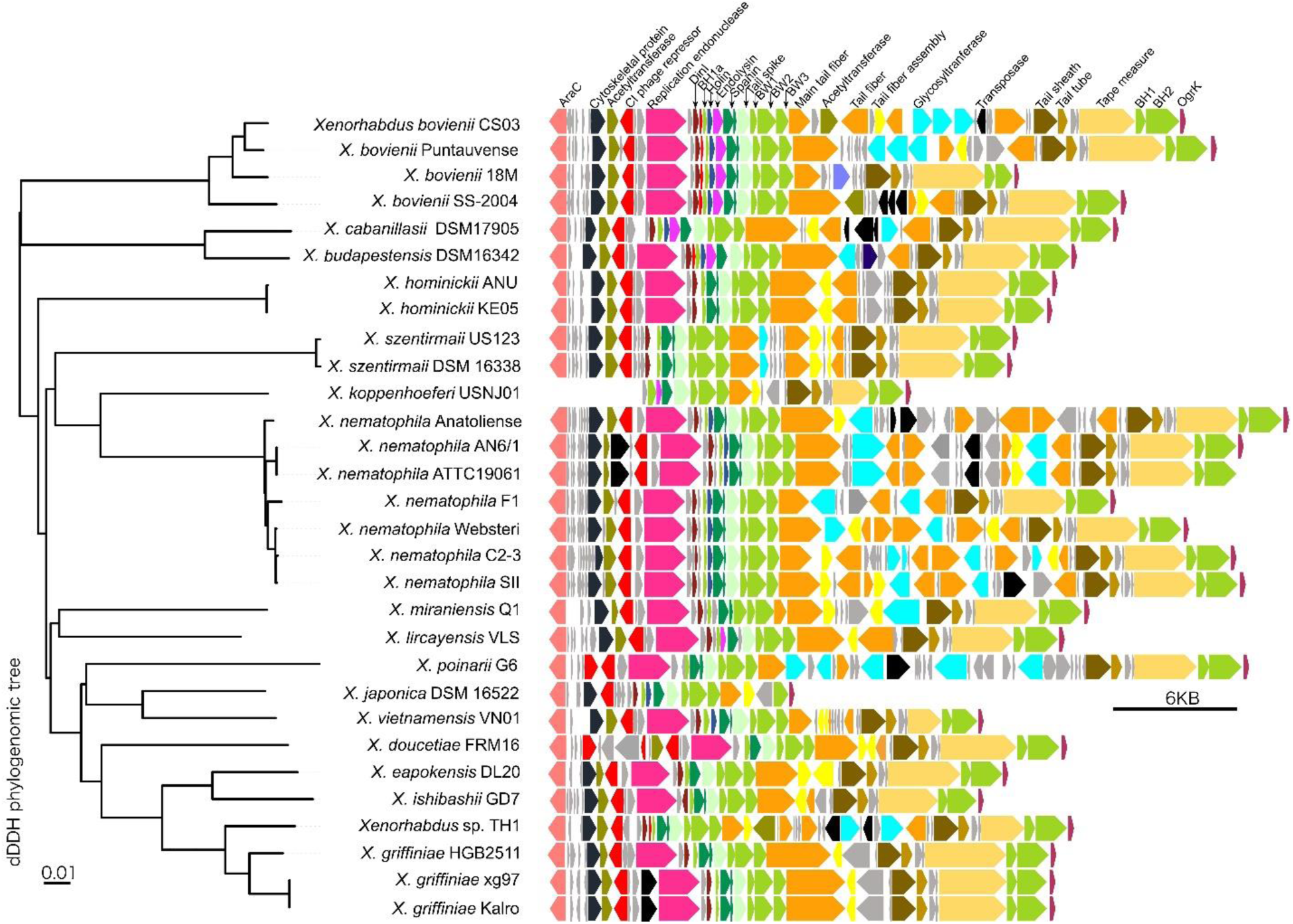
The xenorhabdicin encoding locus contains both conserved and variable regions. An alignment of this locus in 30 strains of *Xenorhabdus* coupled to a phylogenomic distance tree based on digital DNA-DNA hybridization (dDDH) distances. Apart from those of *X. nematophila* FI, Websteri, C2-3, and SII, all shown genome loci were found on one contig and were not split by putative transposition events. The names of the main proteins which are predicted to be encoded by the genes are in the topmost row. Grey indicates that the gene encodes a protein whose predicted function remains unknown. A few loci contained uniquely annotated genes. *X. bovienii* 18M and *X. budapestensis* DSM16342 contained genes that were predicted to code for an anti-CRISPR protein (purple blue) and Mami family restriction endonuclease (dark blue), respectively.

Tapemeasure protein length is positively correlated with tail tube length of bacteriophages (41) and a tailocin (42), and it is possible a similar correlation occurs with xenorhabdicins. Also present in all loci was the *xnpH1* gene that encodes the main tail fiber (XnpH1) (Fig. 2) (20). XnpH1 has a highly conserved *N*-terminal DUF3751 ∼160 amino acid (aa) sequence, which may serve to attach the tail fiber to the baseplate, and a variable *C*-terminal that is predicted to be an RBD that mediates binding to target cell surfaces (21, 43). Indeed, comparison of 97 XnpH1 protein sequences indicated high sequence identity in the *N*-terminal portion, with all 97 proteins appearing nearly identical up to 150 aa, and with just a few exceptions up to 213 aa (Fig. 3). A central region of identity was conserved among some subsets of main tail fiber proteins, with different subgroups sharing different central domains (e.g., amino acids 213-350 are similar among the majority of *X. bovienii* isolates). The dramatic divergence of the 97 main tail fiber *C*-termini, even among conspecific strains (Fig. 3), further supports their predicted function as xenorhabdicin RBDs.

**Figure 3.**
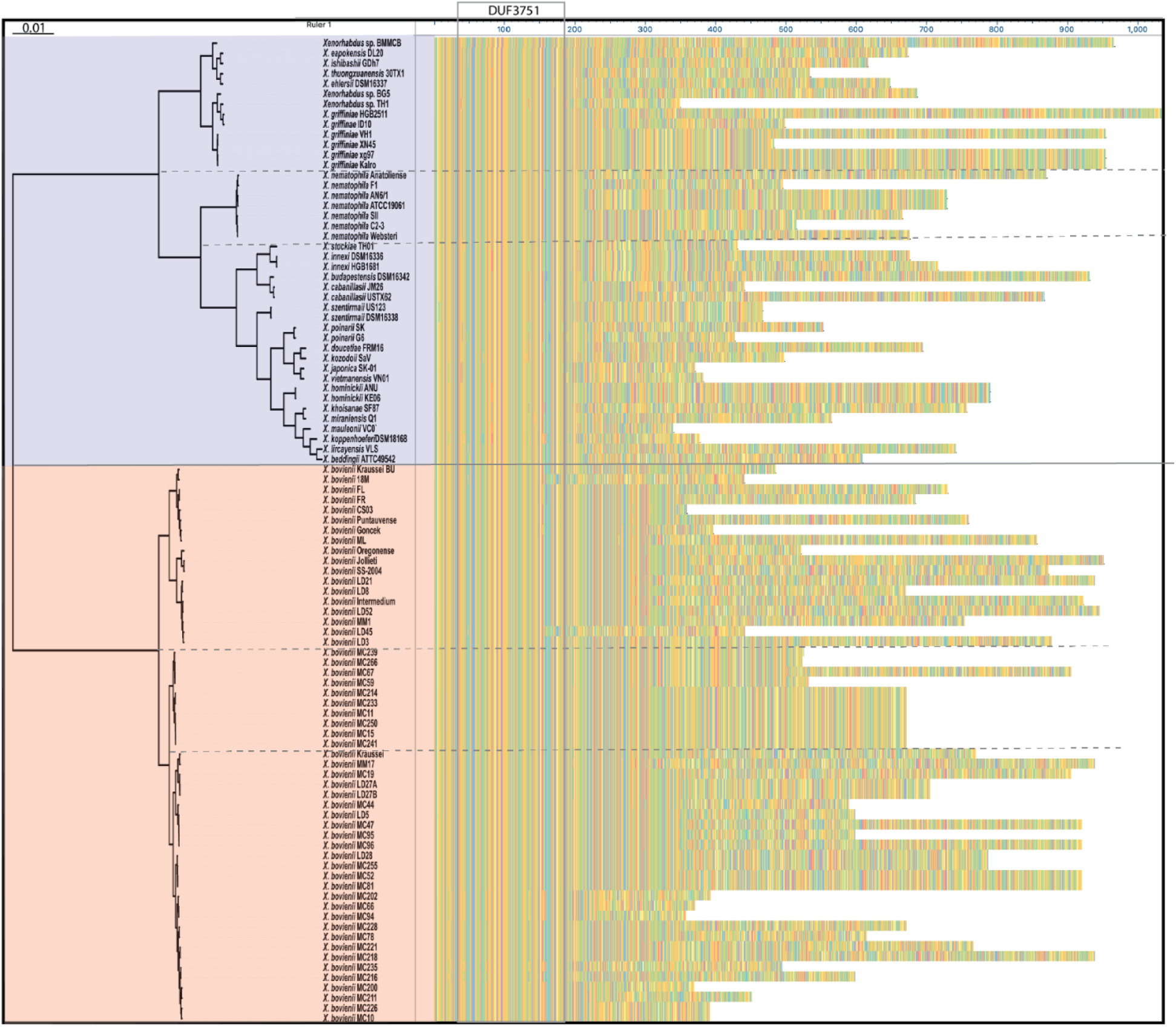
Amino acid sequences of xenorhabdicin main tail fibers (XnpH1) from 97 strains of 27 species from the genus *Xenorhabdus* share a conserved *N*-terminus but have variable middle and *C*-terminal coding regions. On the left is a dendrogram from hierarchical clustering of pairwise Average Nucleotide Identities (ANI) values. The 97 strains fall into two separate clusters, highlighted with blue and pink shading, respectively. Shown on the right are the XnpH1 sequences aligned using DNAStar Megalign software to show regions of amino acid similarity and diversity. The color scheme represents amino acid side chain chemistry: aromatic = yellow; acidic = red; basic = blue; nonpolar = brown; polar = green. The *N*-terminal coding region is highly conserved across all 97 XnpH1 and comprises the ∼160 amino acid DUF3751, which is predicted to function in attaching the tail fiber to the baseplate of the tailocin. The *C*-terminal regions are predicted to contain receptor binding domains (RBDs) that mediate attachment to target cell surfaces. A high level of variability is observed in these RBDs, even among conspecific strains, that in turn likely reflects variability in specificity for target cell surfaces.

### *X. nematophila* tailocins differ in both length and intraspecies killing spectrum

Having observed intra-species variability of the *C*-terminal sequence of the main tail fiber (Fig. 3) we tested whether six strains of *X. nematophila*, ATCC19061, AN6/1, C2-3, and F1, Anatoliense and Websteri (Fig. 3, Table S1), differed both in the tailocins they produced and their concomitant killing spectra. All six *X. nematophila* strains produced visible tailocin structures in extended, contracted, or empty states (except for AN6/1, for which contracted tailocin particles could not be found among sampled images). In some images, the baseplate and tail fibers could be seen clearly (Fig. 4). No other phage-like elements were detected in the preparations (Fig. 4). The tailocin extended structure length, from baseplate to collar, ranged from averages of 171.1nm to 286.6 nm (Fig. 5). These lengths did not correlate with those of predicted *xnp1* locus-encoded tapemeasure protein sequences (Table S2).

**Figure 4.**
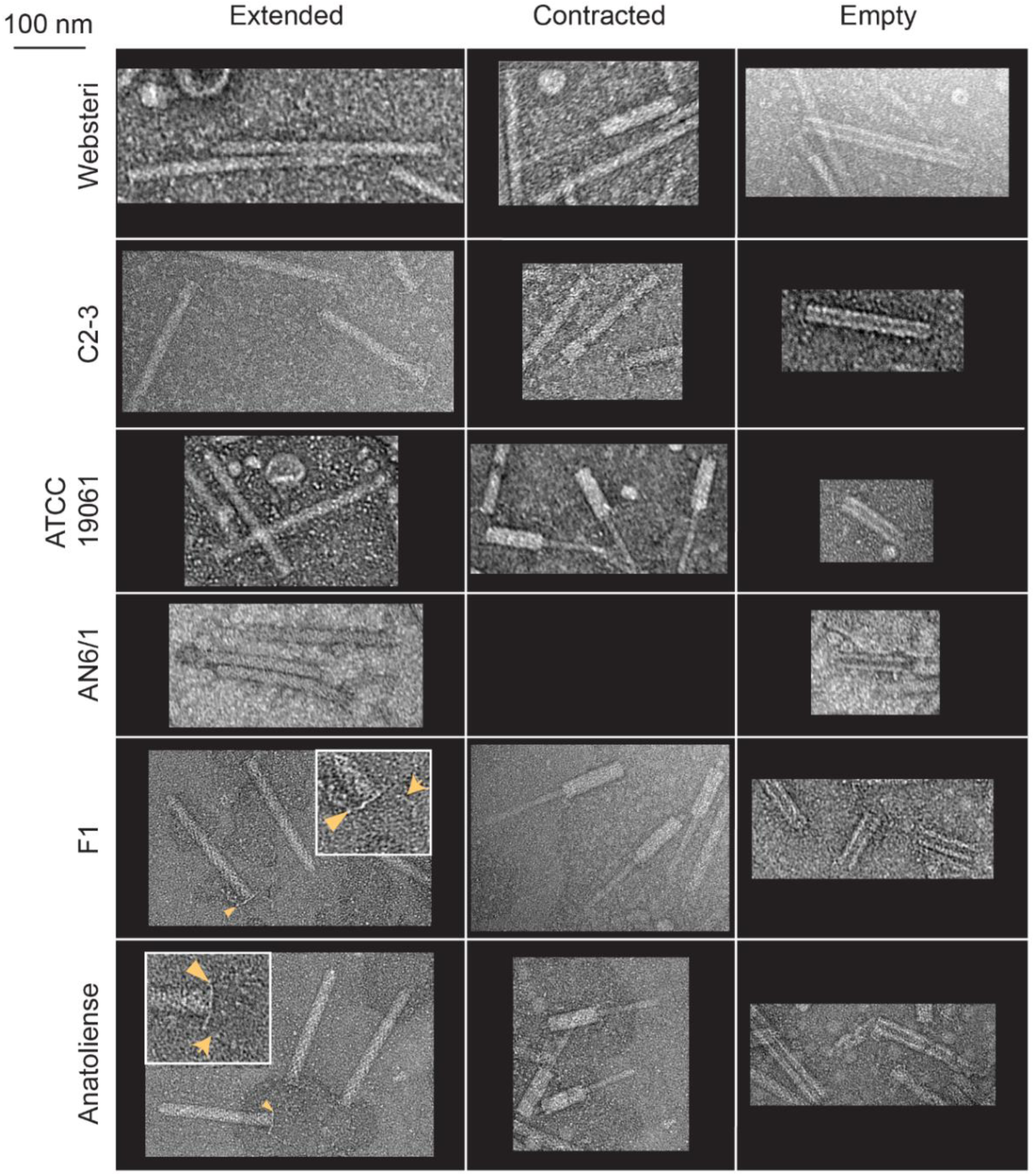
Tailocins are produced by *Xenorhabdus nematophila* strains. Mitomycin C was used to induce tailocin production by six strains of *X. nematophila:* Websteri, C2-3, ATCC19061, AN6/1, F1, and Anatoliense. Pure tailocin preparations were visualized using Transmission Electron Microscopy (TEM), with extended (pre-firing), contracted (post-firing), and empty sheath structures apparent (except for AN6/1 for which no contracted forms were readily apparent in any images). Representative images are shown. In some extended tailocin structures, the baseplate (orange arrowhead) and tail fibers (orange line arrow) were visible. All structures are shown at the same scale (scale bar 100 nm) except for closeup images of Anatoliense and F1, provided to highlight baseplate and tail fiber structures.

**Figure 5.**
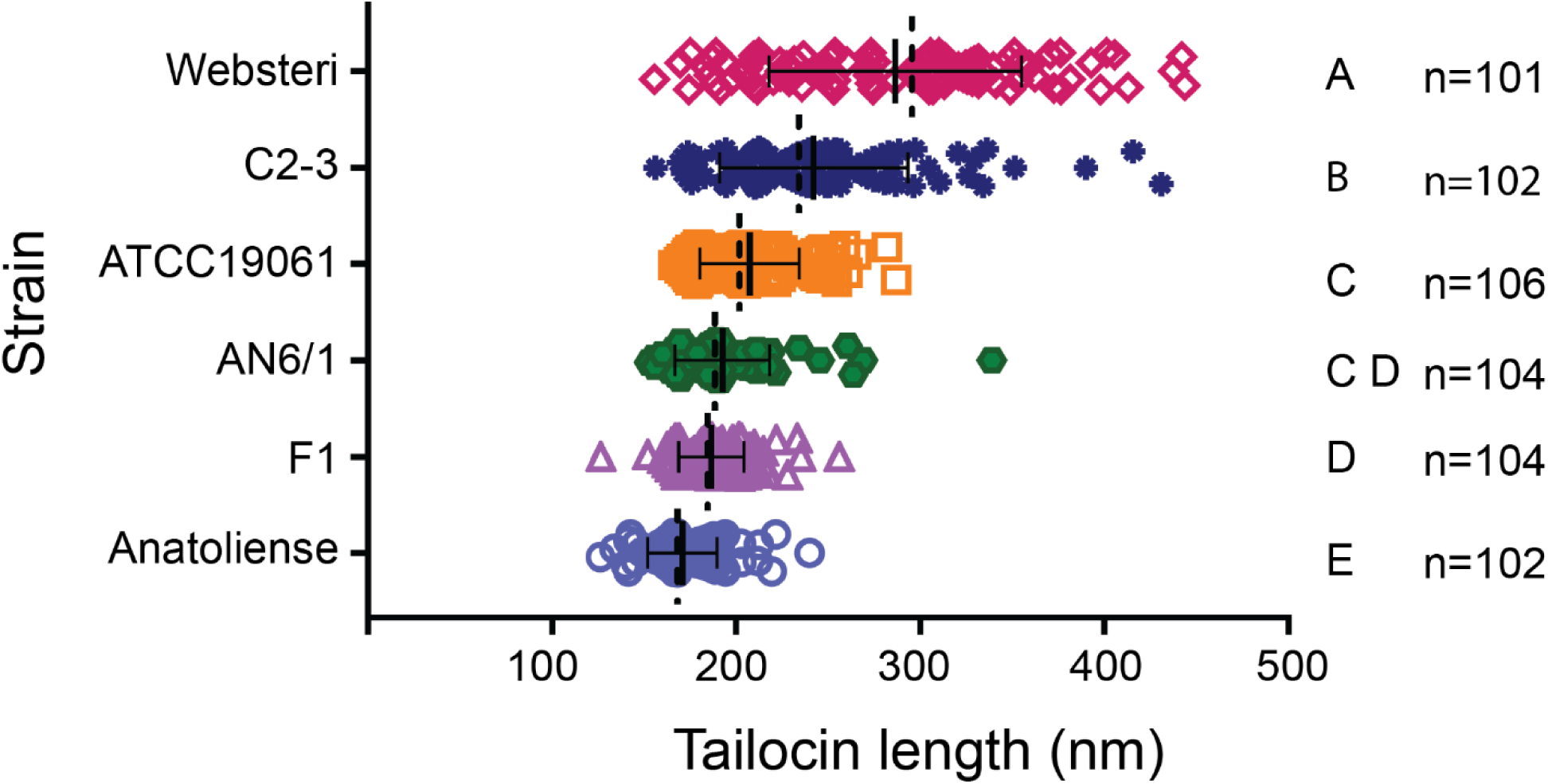
Six *Xenorhabdus nematophila* strains produced tailocins that vary in length. Transmission electron microscope (TEM) images were used to measure the length of extended-form tailocin particles (n ≥ 101) from the tailocin baseplate to collar. Differences in length were observed and these were significant (indicated by the letters to the right) as determined by one way ANOVA multiple comparisons tests. Websteri and Anatoliense strains produced tailocin particles that were significantly longer and shorter, respectively, than those from other strains.

Each actor strain tailocin preparation was assayed for its level of inhibition against each of the other six *X. nematophila* strains. The Anatoliense actor strain exhibited the highest level and broadest spectrum of killing activity, while Websteri and F1 tailocins had little to no inhibitory activity against the other *X. nematophila* strains (Fig. 6A; Supplementary File 2). The closely related strains ATCC19061 and AN6/1 had similar profiles. As actor strains they were strong inhibitors of Anatoliense and to a lesser extent F1 and they were the only recipient strains that were strongly inhibited by actor strain C2-3. *X. nematophila* C2-3 and Websteri were the least susceptible to growth inhibition by any other actor strains. All tailocin preparations inhibited growth of *Photorhabdus luminescens* TT01, a bacterium from a closely related genus, though C2-3 and Websteri showed variability in activity among replicates (Fig. 6A; Supplementary File 2).

**Figure 6.**
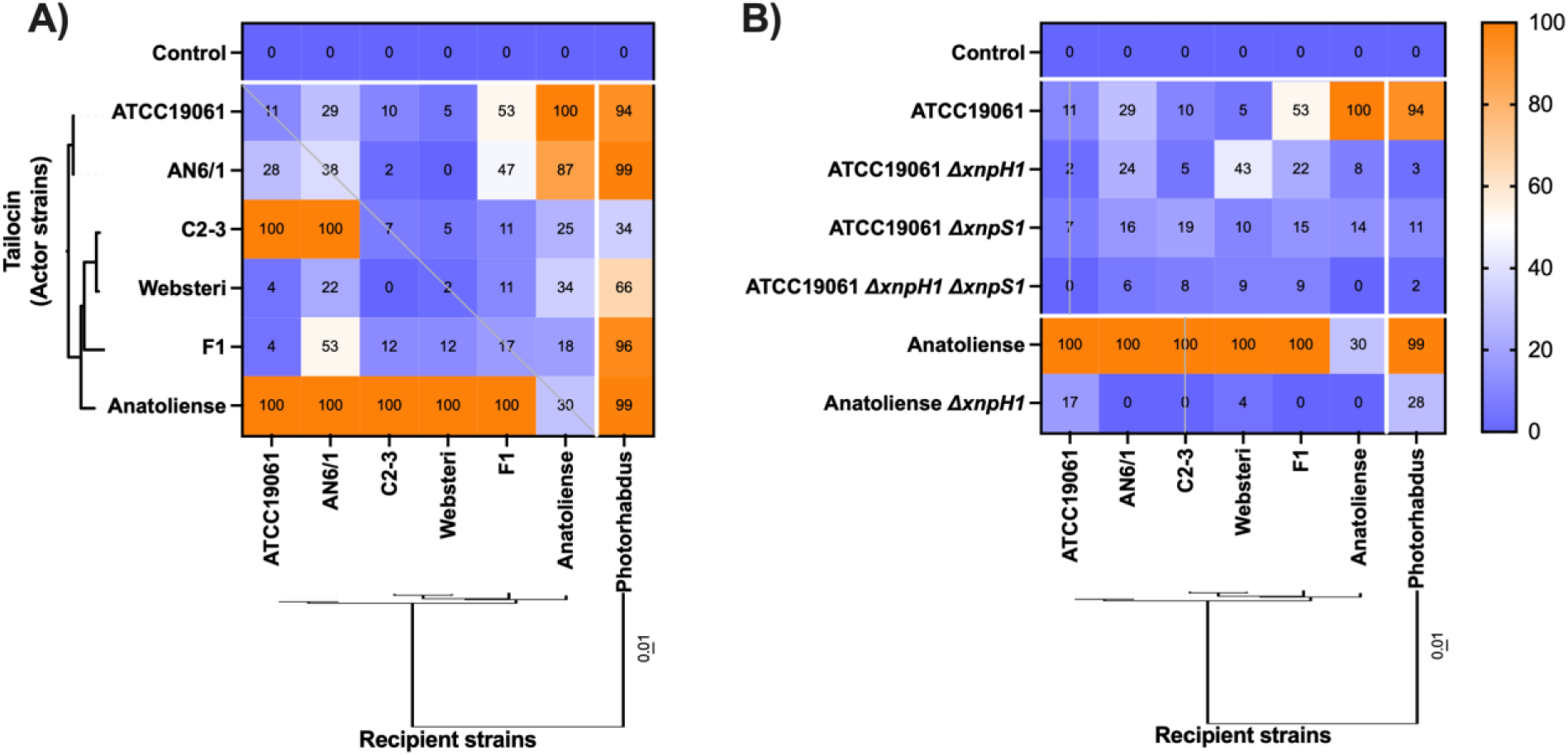
*Xenorhabdus nematophila* xenorhabdicin tailocins differentially inhibit *X. nematophila* recipient strains. Tailocins from each actor strain were tested for inhibition of each recipient strain in a LB liquid killing assay. Tailocin preparations were obtained from wild type *X. nematophila* strains **(A)** or from *X. nematophila* ATCC19061 or Anatoliense mutants in which the *xnpH1, xnpS1*, or both had been deleted **(B)** (Y axis). ATCC19061 and Anatoliense actor strain data are identical in both panels. Tailocin activity was measured against each of the recipient strains (X-axis) and represented as percentage growth inhibition when compared to the LB only control (Supplementary File 2). The heatmap is coupled to a digital DNA-DNA hybridization (dDDH) phylogenomic distance tree that was rooted at midpoint. A light gray line indicates activity assays of actor and recipient from the same strain background.

The dependence of killing activity on an intact *xnp1* locus was tested by creating *xnpH1* (main tail fiber) deletion mutants in *X. nematophila* ATCC19061 and *X. nematophila* Anatoliense. In addition, we created *xnpS1* (sheath protein) and *xnpH1/xnpS1* double mutants in *X. nematophila* ATCC19061. Consistent with previous findings (20), mutants lacking *xnpH1* or *xnpS1* or both were no longer able to kill other stains, indicating that the *xnp1* locus encodes the observed tailocin killing activity (Fig. 6B; Supplementary File 2).

### Sympatric isolates of *X. bovienii* show distinct killing and susceptibility profiles

We next investigated the ecological role of xenorhabdicin tailocins and whether other phage related structures are involved in intraspecies killing. We used a set of 42 *X. bovienii* (Fig. 7A) that had been previously isolated from neighboring forest communities (24), obtained their mitomycin-induced lysates (MILs), and quantified the growth inhibition caused by each MIL against *Xenorhabdus* strains isolated from these communities (Fig. 7B). We then clustered these strains into actor groups based on whether their MILs generally killed the same recipients. This resulted in three actor groups: A1, A2, and A3 (Fig. 7B). We similarly clustered each strain based on whether their growth was inhibited by the same MILs and to what extent, resulting in three recipient groups: R1, R2, and R3 (Fig. 7B). Notably, none of the actor or recipient groups can be considered monophyletic as can be seen by their mapping on the phylogeny (Fig. 7A).

**Figure 7.**
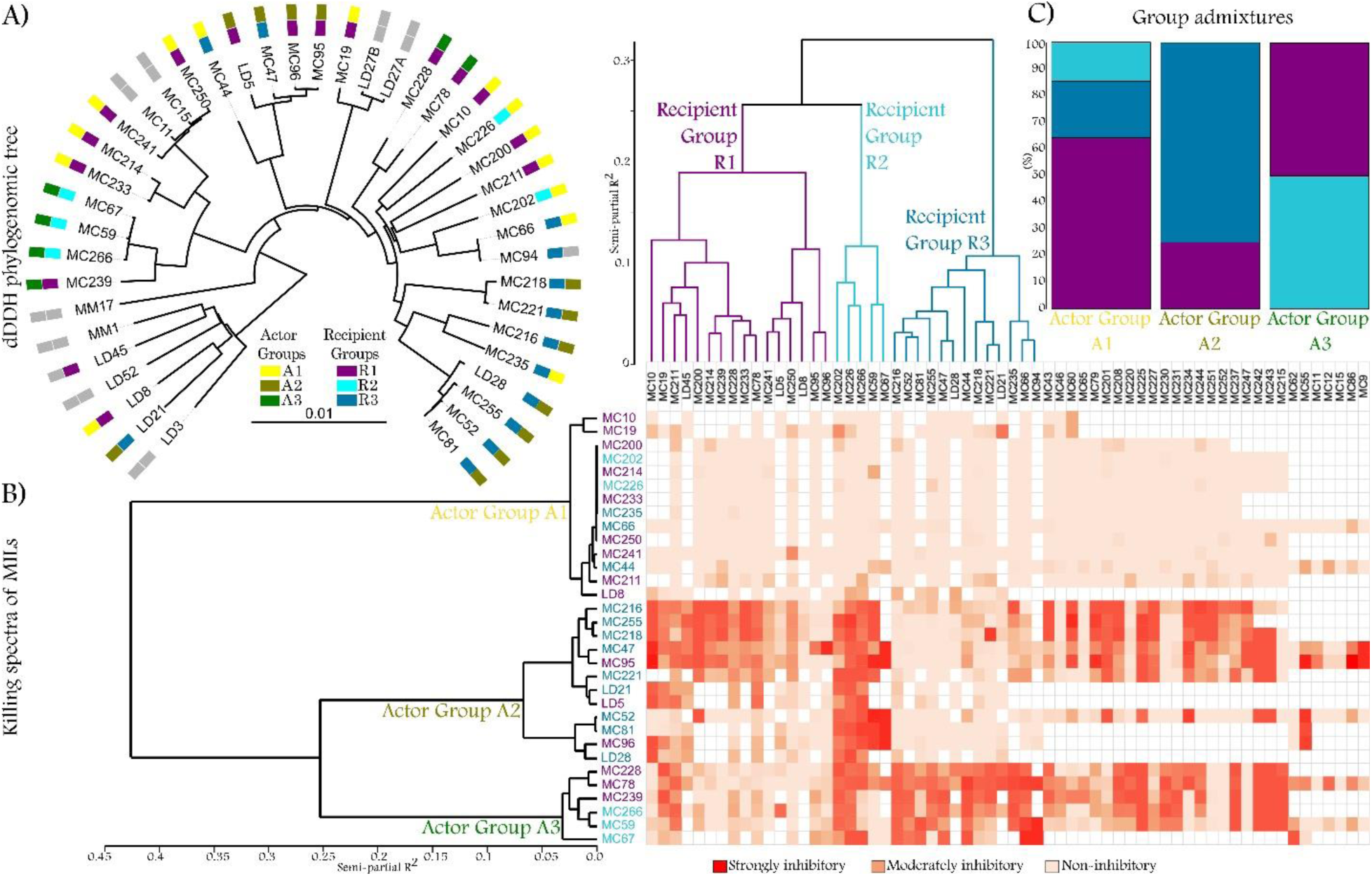
Relationships between the phylogeny of 42 *Xenorhabdus bovienii* strains, the killing spectra of their mitomycin-induced lysates (MILs) and their susceptibility to each other’s MILs. **A)** Phylogeny of 42 strains of *X. bovienii* that were isolated from soil biota from three forest sites (MC, MM, LD) in Indiana. For each strain, actor and recipient phenotypes are mapped as color coded, a grey box indicates that the strain was not phenotyped. **B)** Killing spectrum of each *X. bovienii* MIL. The leftmost column contains names of *X. bovienii* strains from which MILs were produced, clustered based on their actor phenotype. The topmost row contains the names of *Xenorhabudus* strains against which inhibition assays were conducted, clustered based on their recipient phenotype. Along a row, the color of the cell indicates the level to which the MIL produced by strain in the leftmost column, inhibited the strain in the topmost row. Only strains with whole-genome sequences were clustered. **C)** The strains that constituted each of the actor groups belonged to multiple recipient groups.

Examining the killing matrix, A2 and A3 differentiate largely on the ability of their MILs to kill R3 strains. A2 strains are mostly also R3 strains and for the most part their MILs were non-inhibitory to R3 strains. Conversely, A3 MILs are highly inhibitory to R3 strains and none of them are clustered into that recipient group (Fig. 7C). R1 strains are more likely to be killed by MILs from A2 than A3 strains, while R2 strains can be killed by MILs from both. Overall, actor and recipient groups demonstrated high dissimilarity. Their clustering dendrograms (Fig. 7B) had a normalized Robinson-Foulds distance (nRF) between them of 0.93 with 53% branch congruence (BC). Moreover, the actor group dendrogram mapped poorly onto the phylogeny (Fig. 7A, nRF 0.97, BC 3%) as did the recipient group dendrogram (nRF 0.94, BC 6%).

### Both tailocin and phage genes underpin the killing of kin among *X. bovienii*

To identify genes significantly associated with the killing spectra of MILs, we identified GCs that are uniquely found in one of the three actor groups (Fig. 8A). All genomes of strains from actor group A2 contained two GCs that were absent from the other genomes, and this association was the most significant (Fig. 8A). These two GCs coded for a glycosyltransferase and a mannosyltransferase, and were both found in the *xbp1* highly variable region (Fig. 8B). To increase the sensitivity to viral gene calls (44), we redid the association analysis using only Phanotate-annotated *xbp1* loci and identified an additional seven GCs which had not been previously called and that were significantly associated with A2 genomes (Fig. 8B). All these nine GCs that were significantly associated with A2 genomes were in the same genomic region (Fig. 8B).

**Figure 8.**
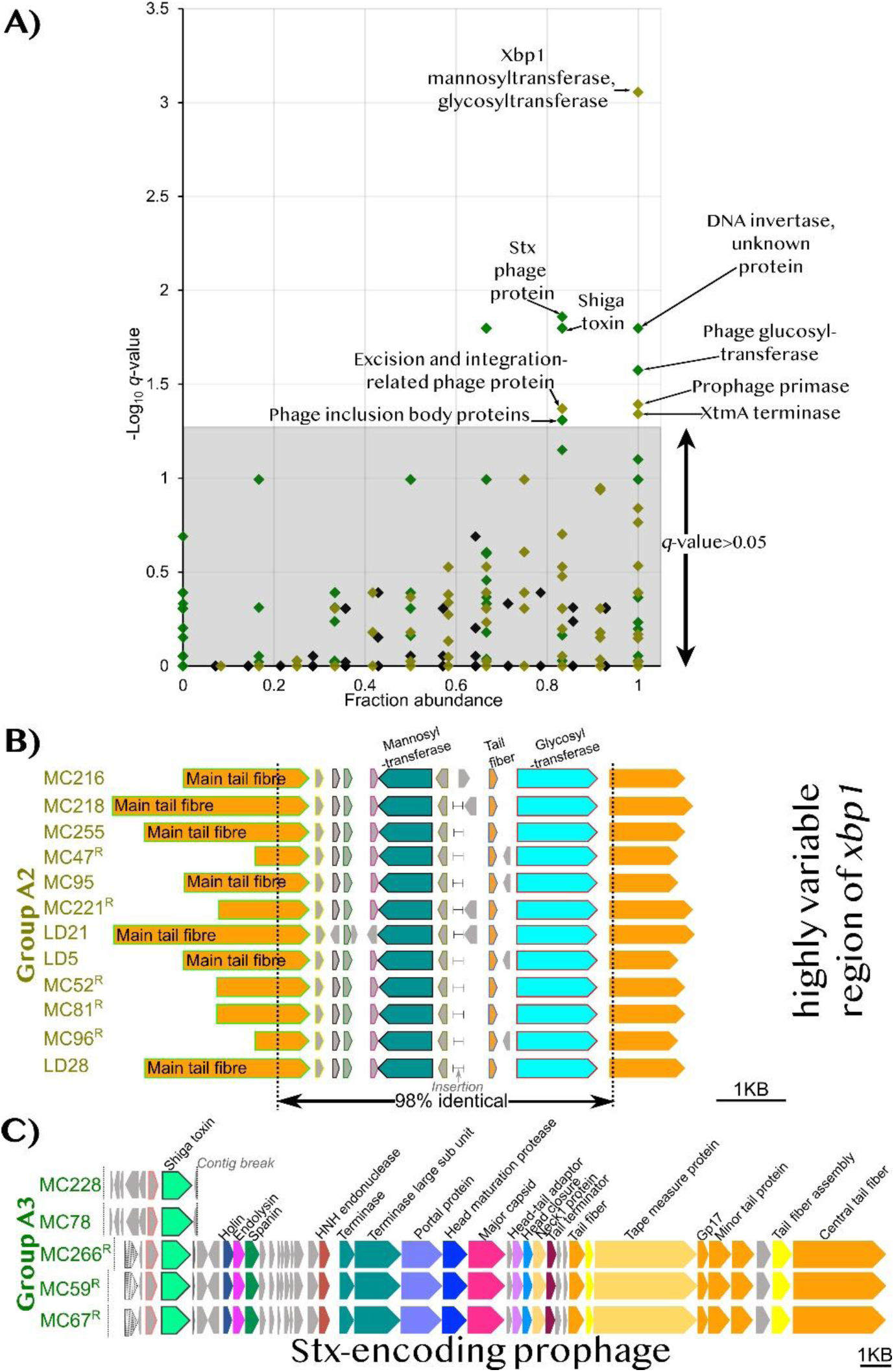
The killing spectra of *Xenorhabdus bovienii* mitomycin-induced lysates (MILs) is significantly associated with genes that encode tailocin and phage proteins. **A)** Significance of genome wide associations between each actor group (A1=black, A2=army green, A3=bright green) and genes against the relative frequency of each gene in an actor group. The proteins predicted to be encoded by the genes are indicated. **B)** Multiple sequence alignment highlighting a 4.6 kilobase (Kb) genomic locus that contained nine genes (in colored outlines) significantly associated with the MIL killing spectrum of group A2. Loci that were inverted to align with the rest are denoted by ^R^ and indicate a likely inversion of the 4.6 Kb fragment in these strains. Consistent with this inversion, the left border of the 4.6 Kb locus was flanked by an invertible repeat sequence AAAGGAA whose palindrome was found in downstream tail fiber genes. The strain MC216 locus contained a 173-base pair (bp) insertion whose position is indicated by a gap in the other sequences. **C)** An alignment of Stx-encoding prophage that contained a gene predicted to encode the Shiga toxin (Stx) which was significantly associated with the killing spectra of group A3. An unknown gene (pink outline) directly upstream of *stx* was similarly associated with A3 killing spectra. The gene at the start of the MC266, MC59 and MC67 loci and denoted by wavy lines was predicted to encode an anti-terminator protein Q.

Multiple sequence alignment of the genome region containing all these nine GCs revealed a 4.6 Kb fragment that had a percentage pairwise identity of 98% (Fig. 8B) for the 12 A2 genomes, notwithstanding its location in the highly variable *xbp1* region which had a percentage pairwise identity of 25% for 42 *X. bovienii* strains of Fig. 7A. For seven A2 genomes, this 4.6 Kb locus contained the main tail fiber RBD-encoding gene fragment (Fig 8B) demonstrating a direct and significant association between RBDs and *Xenorhabdus* MILs killing spectra. For the remaining five A2 genomes (MC47, MC221, MC52, MC81, MC96), the orthologous tail fiber gene fragment was of an extra tail fiber gene. For these five genomes, only when the entire 4.6 Kb fragment was inverted did it align with the seven (Fig. 8B), demonstrating a DNA inversion that is directly associated with tailocin killing spectra.

Like tailocins, phages are possibly exploited by *Xenorhabdus* bacteria to compete against close kin. We found several cargo prophage genes significantly associated with the killing spectra of *X. bovienii* MILs (Fig. 8A). For A3 strains, their MIL killing spectra were associated with two neighboring genes, including one that encodes a Shiga toxin (Stx), found in a prophage (Fig. 8C). Moreover, an unknown prophage (contig JAILTI010000026 of assembly GCA_029104845.1) in A3 genomes contained a glycosyltransferase-encoding gene which was also significantly associated with A3 MIL killing spectra (Fig. 8A). A3 MIL killing spectra were also significantly associated with two neighboring phage inclusion body genes (Fig. 8A) that were in another unknown prophage (141,643-150,837 on contig JAILTI010000007 of GCA_029104845.1). A DNA invertase gene was significantly associated with A3 MIL killing spectra (Fig. 8A).

In A2 genomes, three different prophages each ferried genes that were significantly associated with A2 MIL killing spectra. First was an unknown prophage (locus 110,842-146,771 on contig JAILSP010000008 of assembly GCA_029104565.1) that contained a gene predicted to encode an excision and integration-related phage protein (Fig. 8A). Second was the prophage of contig JAILSP010000028 on assembly GCA_029104845.1 that contained a significantly associated *xtmA* gene (Fig. 8A). Third was a prophage at 115,465-146,771 on contig JAILSP010000008 of assembly GCA_029104565.1 that contained a gene predicted to code for a DNA primase that was significantly associated with A2 MIL killing spectra. Together, the significant associations between intraspecific inhibitions and phage and tailocin genes indicate that tailocins, genomic rearrangements, and bacteriophages are important to the functional diversity of interactions among *Xenorhabdus* bacteria.

### The susceptibility of *X. bovienii* strains to death by MILs is associated with genes that encode the biosynthesis of O-antigens and *Photorhabdus* virulence cassettes

A variety of genes underpin the susceptibility of an *X. bovienii* strain to inhibition by MILs since we identified numerous GCs significantly associated only with either R2 or R3 genomes (no GCs were significantly associated only with R1 genomes, Fig. 9A). For R2 and R3 genomes, ten and twelve of these genes, respectively, were colinear, and based on this, we focused our investigations on these two regions (Fig. 9B, C).

**Figure 9.**
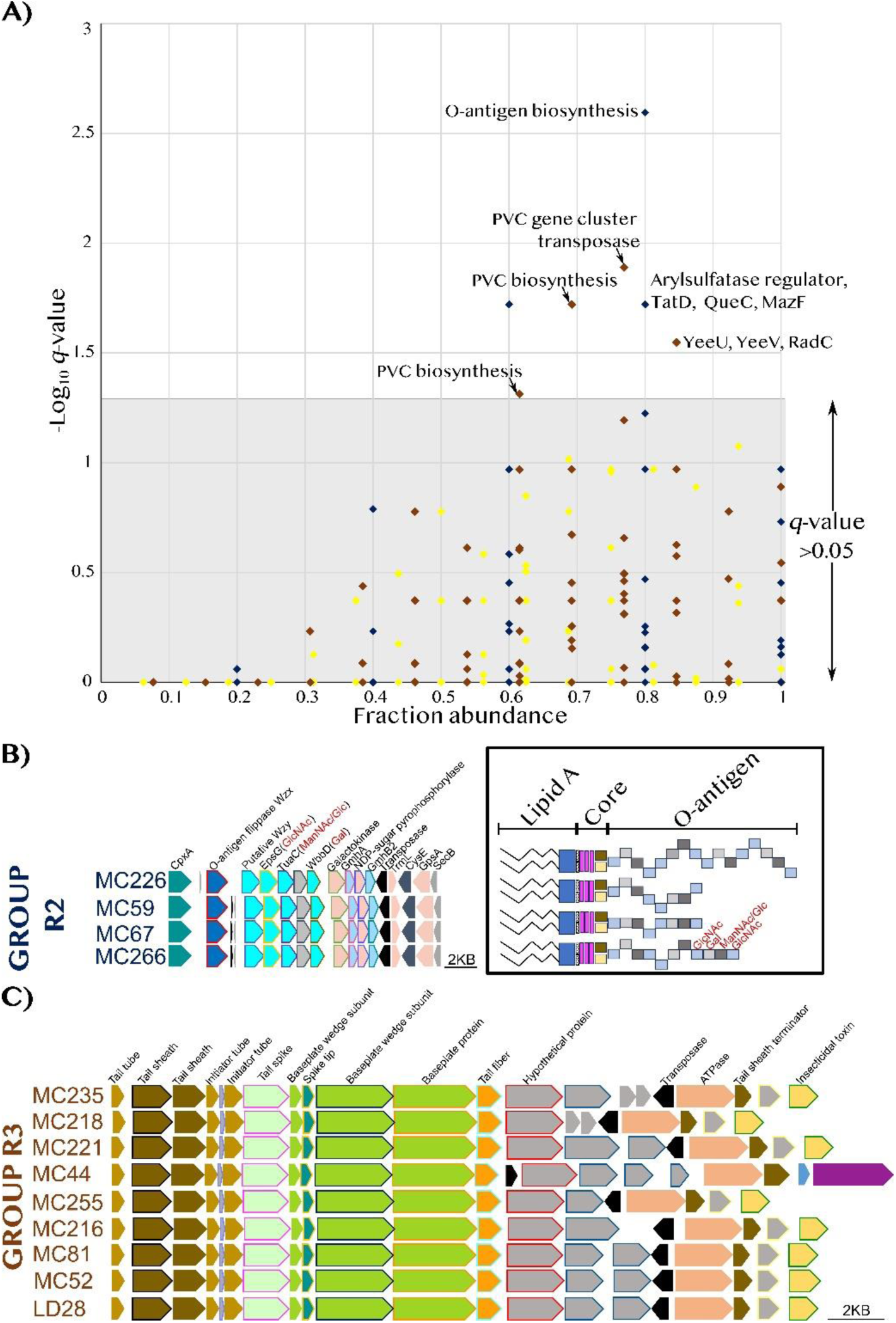
The susceptibility of *Xenorhabdus bovienii* strains to mitomycin-induced lysates (MILs) from close relatives is most strongly associated with genome loci that encode O-antigen biosynthesis and production of *Photorhabdus* virulence cassettes (PVCs). **A)** Significance of genome wide associations for each recipient group (R1=yellow, R2=blue, R3=brown) versus the relative frequency of each gene in a recipient group. **B)** A schematic of genome loci that encode O-antigen biosynthesis in group R2 strains a model of corresponding O-antigen structure. **C)** Genome loci that encode PVC biosynthesis in group R3 strains. The gene clusters, a set of orthologous genes, that were specifically enriched are outlined and genes with the same color outline in a locus, belong to the same gene cluster. The proteins predicted to be encoded by genes are written on the top rows.

The LPS of R2 strains likely contains a unique *O-*antigen that underpinned their increased susceptibility to A2 and A3 MILs, since 4/5 of their genomes contained 10 GCs all found in a genomic region that encodes *O*-antigen biosynthesis (45) (Fig. 9B). This putative R2-specific *O-*antigen likely contains glycan residues of galactose, *N*-acetylglucosamine and *N*-acetyl-*D*-mannosamine, as the corresponding locus contained genes predicted to encode WbbD, EpsG and TuaC (46–48). Interestingly, the genome of MC202, the R2 strain that lacked orthologs to these 10 GCs encoded UDP-galactopyranose mutase (Supplementary File 1) the central enzyme in the biosynthesis of the non-mammalian monosaccharide *D*-galactofuranose that is also an *O*-antigen glycan residue (49). Altogether, these findings indicate that *Xenorhabdus O*-antigen glycan residues are the likely molecular receptors of bacteriophages and tailocins in MILs.

Most R3 strains encode “*Photorhabdus* virulence cassette” (PVC) biosynthesis, and this is significantly associated with R3 susceptibility to A3 MILs and resistance to A2 MILs (Fig. 7B). Specifically, 9/13 R3 genomes harbored genes that coded for PVC tail sheath, tail spike, spike tip, baseplate wedge subunit, baseplate protein, tail fiber, three unknown PVC proteins and an insecticidal toxin (Fig. 9B). This indicates possible links among PVCs, tailocins and bacteriophages in mediating reciprocal competitive effects.

## DISCUSSION

The simultaneous association of *Xenorhabdus* with two eukaryotic hosts fosters a high level of competition among strains, making these bacteria valuable systems for evaluating the effect of conspecific competition on genome diversification. Our comparative analysis of 97 *Xenorhabdus* genomes revealed that they have among the highest proportion of phage-related genes found in bacterial genomes These genes underlie the striking variation in proteome sizes across the *Xenorhabdus*, with 9.8% of protein-coding genes found in phage-associated regions. The association of non-cargo phage genes with proteome size suggests that phages are the primary porters used by *Xenorhabdus* genomes to acquire genes and that phage-associated regions can facilitate recombination. Our pangenome findings contradict the conclusion based on the genus *Arsenophonus* that loss of CRISPR systems is a general prerequisite for this phage-mediated genome expansion upon endosymbiosis (50), since except for that of *X. stockiae* TH01, all *Xenorhabdus* genomes encoded components of CRISPR systems, corroborating prior analyses of CRISPR systems from *X. griffiniae* and *X. nematophila* (51, 52). Perhaps the retention of CRISPR in *Xenorhabdus* is linked to its potential role in associating with the nematode during transmission between insects (52). Indeed, across a broader range of bacteria, prophage content is only reduced if CRISPR systems have many spacers, indicating active functioning in phage defense (53). Consistent with this prior work, *Xenorhabdus* fits the pattern of higher prophage content in faster growing and pathogenic bacteria, as the boom-bust cycles concomitant with these lifestyles is thought to favor phage lysogeny (53, 54).

Although no active phages have been described infecting the *Xenorhabdus* genus, the phage-like tailocin, xenorhabdicin, is an important mediator of competition, capable of protecting an insect cadaver from a related entomopathogenic bacterium, *P. luminescens* (20, 21). The adaptive importance of genomic phage-derived regions has been well established in bacterial pathogens as many virulence factors and biological weapons used for host colonization are encoded by phage regions (55). Here, we show the importance of such regions in intraspecific competitive interactions. All genomes we examined encoded xenorhabdicin tailocins, indicating an ancestral origin of the locus for the genus that has evolved to range across strains more than three-fold in size. The diversity found in this genomic region across strains, even within a single species, suggests that it is under selection. Indeed, the phenotypic diversity among both *X. nematophila* tailocin size and killing activity spectrum indicates that these competitive interactions are a selective force driving xenorhabdicin diversification. In addition, the hypervariability of xenorhabdicin main tail fiber *C*-terminal domain amino acid sequences corroborates its predicted function as a diversifying receptor binding domain (21).

While aspects of *xnp1* locus sequence divergence do not impact the killing ability of six *X. nematophila* xenorhabdicins against a common enemy, *P. luminescens*, it does impact xenorhabdicin particle size and killing activity within the species. The ecological importance of this diversification is highlighted by our finding that there are significant associations between both xenorhabdicin and prophage genes and intraspecific variation in killing profiles of 42 *X. bovienii* sympatric strains. These data suggest that despite sequence divergence, xenorhabdicin loci preserve overall xenorhabdicin effectiveness against distantly related bacteria, including *P. luminescens* that pose a threat to the overall symbiosis between *Xenorhabdus* and its *Steinernema* nematode host (20). Intriguingly, our data in both *X. nematophila* and *X. bovienii* demonstrate that xenorhabdicin sequence divergence expands its functionality as a precision weapon against other *Xenorhabdus* species. Since strains of the same species associate with the same nematode host, they are direct competitors with each other for host colonization. This colonization is required for transmission to a new insect host, and variant strains of the same species therefore pose a significant threat to each other for fitness.

Unlike loci that encode similar rigid contractile type tailocins in *Pseudomonas* (9, 56, 57), *Burkholderia* (58), *Clostridium* (42) and *Dickeya dadantii* (59) and like those of *Budviciaceae* (60), xenorhabdicin-encoding loci encode a large diversity of receptor binding domains, since they contained up to six extra tail fiber genes (Fig. 2). These extra tail fiber genes lack the conserved *N*-terminal domain of the main tail fiber that is expected to mediate attachment to the tailocin baseplate yet still encode receptor binding domains (21). For five *X. bovienii* strains, extra tail fiber genes were significantly associated with killing spectra of putative xenorhabdicins contained in MILs. In these five strains, an extra tail fiber gene associated with xenorhabdicin killing spectra was inverted to replace the *C*-terminal encoding fragment of their own main tail fiber gene, resulting in their loci matching those of the other members of this phenogroup. This suggests that extra tail fiber genes are exchanged with the *C*-terminal encoding fragment of the main tail fiber gene, probably through site-specific recombination (61, 62), to alter killing spectra of xenorhabdicins. Site specific recombination requires a recombinase and invertible repeat sequences that flank the DNA fragment to be inverted (63). This type of recombination alters killing spectra of tailocins from *Pectobacterium carotovorum* (61) and is suggested to do the same for *Pantoea stewartii* tailocins (62), whose encoding loci are strikingly similar in gene collinearity to those which encode xenorhabdicin. A major difference between the two types of loci is that xenorhabdicin-encoding loci did not contain predicted DNA invertase genes. However, all 97 genomes did contain genes in other loci that were predicted to encode various site-specific recombinases, including a DNA invertase gene that was significantly associated with the killing spectra of A3 strains. These data indicate that DNA inversion within xenorhabdicin-encoding loci may be mediated by multi-site recombinases (63). We speculate that xenorhabdicins from a single cell whose genome, for example, encodes four xenorhabdicin receptor binding domains will release xenorhabdicin particles that differ by having one of four tail fibers. Such mixed tail fiber tailocin preparations have been experimentally demonstrated for *Pseudomonas chlororaphis* tailocins (56). We found that *X. nematophila* xenorhabdicins clonally vary in actual baseplate-to-collar length and this wsa unexpectedly not associated with tape measure gene length (42). Possibly, there may be an association between the receptor binding domain identity and tailocin length due to mechanical constraints.

Like tail fiber genes, glycosyltransferase genes were frequently found in the hypervariable region of the xenorhabdicin-encoding locus and were significantly associated with the killing spectra of putative xenorhabdicins and phages in MILs from *X. bovienii* strains. Bacterial glycosyltransferases catalyze cell envelope biogenesis, and when encoded by prophages, they are used to tailor the LPS of a bacterium to make it nonspecific to the receptor binding domains of corresponding bacteriophages released by sister cells (64). Hence, we propose that the mannosyltransferases and glycosyltransferase genes associated with the killing spectra of putative xenorhabdicins in MILs from A2 strains mediate a “self-resistance mechanism” by encoding modifications to the LPS making it non-specific to the receptor binding domains of xenorhabdicins released by a sister cell. This hypothesis would not only explain the preponderance of glycosyltransferases genes in xenorhabdicin-encoding loci but also contradict the suggestion that tailocin-encoding loci do not contain genes encoding self-immunity (65). The prophage-ferried glycosyltransferase gene that was significantly associated with killing spectra of putative phages in MILs of A3 strains likely encodes a similar type of self-resistance albeit to the corresponding bacteriophage.

Through the *X. bovienii* genome-wide-association study, we found two loci that were significantly associated with susceptibility profiles. One was a significant association between genes from the O-antigen biosynthesis locus of R2 strains and their susceptibility to inhibition by putative phages and xenorhabdicins. This finding is consistent with those from *Pseudomonas syringae, P. fluorescens* and *P. protegens* where either mutations and gene disruptions in or entire deletions of O-antigen biosynthesis loci reduced the sensitivity of mutants to tailocin killing (12, 65, 66). Moreover, among 32 *P. aeruginosa* strains, sensitivity to death by tailocins was correlated with serotypes determined by O-antigens (67). Another association with susceptibility was with PVC biosynthesis genes, which could be due to common transcription regulation of tailocin, phage, and PVC encoding loci. We speculate that when PVC biosynthesis transcription is activated, that of the xenorhabdicin hypervariable region is repressed. The latter would prevent LPS alteration by glycosyltransferases encoded by the *xbp1* locus and hence alter the strain’s susceptibility to various xenorhabdicins.

In conclusion, the high phage gene content of *Xenorhabdus* genomes sets the stage for complex interactions among phage-encoded killing mechanisms. Competitive inhibition between *Xenorhabdus* strains is underpinned by one population releasing either tailocins or phages with receptor binding domains that bind to specific glycan residues on the cell surface of the inhibited population. These competitive interactions drive diversification of both receptor and phage and tailocin loci, as strains battle to gain dominance through production of novel weapons and defense. *Xenorhabdus* are an attractive system to decipher the enigmatic code by which each receptor binding domain sequence targets specific target cell surface glycan residues. This deciphered code would establish the development of bespoke, safe and effective (57), antimicrobials for treatment of diseases caused by not only bacterial but also protozoan pathogens, since a few of the latter have cell surfaces decorated with the same *D*-galactofuranose residue that is a putative receptor targeted by *Xenorhabdus* tailocins and phages.

## MATERIALS AND METHODS

### Comparative Genome Analyses

To determine the proportion of mobile genetic elements, their composition, and genomic loci within *Xenorhabdus* genomes, we constructed a pangenome of the genus from 97 publicly available genomes, whose accession numbers are listed in Supplementary File 1. We used Anvi’o 8 (68) for pangenome construction as we had previously described (51) to create a pangenome with the following crucial parameters: Markov clustering (MCL) inflation of two; use of Diamond (69) to calculate amino acid (aa) sequence similarities and exclusion of partial gene calls. To compute average nucleotide identities (ANI) within anvio we used fastANI (70) created a dendrogram from pairwise ANI values by hierarchical clustering, using the Euclidean distance metric and Ward Linkage method. We obtained a spreadsheet of the pangenome (Supplementary File 1) and used it to calculate the number of genes, gene clusters (GCs), and their distribution in the pangenome’s core, accessory and strain-specific regions.

For the 97 genomes, we determined any significant associations between mobilome and proteome size by calculating the Pearson’s correlation coefficient between size of the mobilome in a genome and 1) genome length 2) total protein-coding genes. Further, we examined the correlation between size of each of the three types of mobilome (non-cargo phage genes/non-cargo plasmid genes/transposase-encoding genes) and the total number of protein-coding genes per genome. Prophage/phage-like loci were identified with Genomad (39). Xenorhabdicin-encoding loci were identified by querying NCBI-annotated genomes for loci flanked by *araC* and *ogrK.* Loci were reannotated using both Bakta (41) and Phannotate (42), visualized in Geneious, and used to create gene diagrams as well as assign predicted functions that were supported by more than one of the annotation programs used. For phylogenomic analyses, digital DNA-DNA hybridization distance trees were reconstructed as previously described (43). Tree comparisons were made using normalized Robinson-Fould distances in ETE3 (44).

Genome wide association analyses were conducted to identify genes that were associated with either with the killing spectra of *X. bovienii* mitomycin induced lysates (MIL) or sensitivity of a bacterium to MILs, using a binomial general linear model, Rao test and calculation of *q*-values from *p*-values as detailed in a previously described workflow (45).

### Bacterial growth conditions

Table S1 lists the bacterial strains and plasmids used in this study. All *Xenorhabdus* strains were cultured in Luria Bertani (LB) media consisting of tryptone (10 g/L), yeast extract (5 g/L), and NaCl (10 g/L), adjusted to pH 7.0 and, when needed, solidified by supplementation with agar (20 g/L). Unless otherwise noted, *X. nematophila* were grown at 30°C and *X. bovienii* were grown at 28°C. *E. coli* strains harboring the pKNGkan vector were grown at 37°C in LB broth supplemented with kanamycin (50 µg/mL). The *E. coli* S-17 λ pir DAP-dependent strain, used to generate the donor strain, was grown at 37°C in LB broth supplemented with diaminopimelic acid (DAP) and kanamycin. *E. coli* transformants were cultured on LB agar plates containing DAP and kanamycin. For the selection of recombinant strains, LB agar (15 g/L) plates supplemented with sucrose (50 g/L) were used. Exconjugants were selected on LB agar plates supplemented with kanamycin (50 µg/mL).

### Construction of *xnpH1* and *xnpS1* deletion mutants

To test whether x*np1* locus genes are necessary for tailocin production by *X. nematophila* ATCC19061 and Anatoliense strains, site-specific homologous recombination was used to delete two core tailocin genes: *xnpH1* (main tail fiber) and *xnpS1* (sheath). We generated single tail fiber mutants in both ATCC19061 (*ΔXNC1_1193)* and Anatoliense (*ΔXNA1_v220011*). Additionally, an *xnpS1* sheath mutant and a *xnpH1 xnpS1* double mutant were created in the ATCC19061 strain background. To generate deletion constructs, primers were designed to amplify ∼1 kb upstream and downstream of each gene to be deleted (Table S2) and amplified products were cloned into the vector pKNGkan.

Amplification primers for upstream and downstream fragments included 20–30 bp overlaps homologous to sequences flanking the pKNGkan vector (Table S2) multiple cloning site, to facilitate fusion using HiFi assembly (NEBuilder® HiFi DNA Assembly). pKNGKan is a derivative of pKNG101 (71) in which the NotI-ApaI fragment containing a streptomycin resistance cassette has been replaced by a ∼1 Kb NotI-ApaI fragment derived from pKanWor containing a Km-resistance-cassette (aminoglycoside phosphotransferase COG3231) (Table S1).

PCR reactions mixes contained 200 μM dNTPs, 0.5 μM of the forward and reverse primers, 0.02 U/μl Q5 High-Fidelity Master Mix (New England Biolabs) and template DNA: 100 ng/μl *Xenorhabdus* genomic DNA (isolated with Invitrogen PureLink^TM^ Genomic DNA Mini Kit) or 20-50 ng/μl plasmid DNA (purified with Wizard^R^ plus SV minipreps DNA Purification systems). The thermocycling conditions were an initial denaturation at 98°C for 3 min, 35 cycles of 98°C for 10 s, annealing for 30 s, extension at 72°C (1 Kb/30 s), followed by a final extension at 72°C for 2 min. The PCR products were then analyzed by agarose gel electrophoresis. To enrich linear, amplified pKNGkan and prevent carryover transformation of the circular template, PCR was first performed on serial dilutions of 100 ng/μl pKNGkan.

The lowest concentration yielding positive PCR results was then subjected to overnight DpnI digestion (8 μl PCR product, 1 μl DpnI [NEB R0176S/L] and 1 μl nuclease-free water) at 37°C, which should cleave cell-derived, methylated, circular, but not amplified, unmethylated, linear DNA. The DpnI enzyme was then inactivated at 80°C for 20 min. All linearized PCR products, including DpnI-digested pKNGKan, were purified, concentrated, and purified using a Zymo DNA Clean & Concentrator Kit. For each deletion construct, upstream genomic fragments, downstream genomic fragments, and linearized pKNGKan were combined in equimolar concentrations with HiFi master mix and incubated at 50°C for 1 h. The resulting mix was transformed into *E. coli* S-17 λ pir dap-dependent cells using the NEB High Efficiency Transformation Protocol (C2987H) V1 and positive transformants were selected for on LB with diaminopimelic acid (DAP) and kanamycin. Briefly, 5 µL of the plasmid construct was added to 50 µL of *E. coli* S-17 λ pir DAP-dependent cells. This mixture was incubated on ice for 30 min and then heat-shocked at 42°C for 30 s. The mixture was further incubated on ice for 3 min. Next, 950 µL of recovery LB medium containing 0.02% diaminopimelic acid (DAP) was added to the cells. The bacterial cells were cultured at 37°C with shaking for 1 h. The transformants were selected for on LB supplemented with 0.02% DAP and kanamycin (50 µg/mL final concentration), since the *E. coli* cells used for this transformation depend on DAP for growth, and the pKNGkan plasmid contains a kanamycin resistance gene. Transformants were assessed by colony PCR using insert-specific primers (forward primer of the upstream sequence and reverse primer of the downstream sequence) (Table S2). Plasmids were extracted from the positive clones and further verified by DNA sequencing (Plasmidsaurus).

*X. nematophila* were transformed with plasmid constructs by conjugation with *E. coli* donor cells. Briefly, 5 mL LB supplemented with 0.02 % DAP and kanamycin (50 µg/mL final concentration) was inoculated with a single colony of the recipient *X. nematophila* ATCC19061 (HGB800) and the donor *E. coli* and incubated overnight at 30 °C with shaking until late-log phase growth (OD_600_ = 0.8). Cells were harvested by centrifugation at 13,000 x ***g*** for 10 min at room temperature, washed twice with phosphate buffered saline (PBS), and resuspended in 500 μL PBS. The donor and the recipient cells were mixed in equal ratios and spotted onto LB plates supplemented with DAP to support the growth of S17-1 λ*pir* DAP-dependent *E. coli* and facilitate transfer of the deletion construct.

The conjugation mixture was incubated overnight at 30 °C, resuspended in LB, and plated onto LB agar supplemented with kanamycin (50 μg/mL final concentration). The absence of DAP eliminated the donor strain, while kanamycin selected for exconjugants (merodiploids) with the deletion vector. Post-conjugation, exconjugants in which the plasmid had integrated into the chromosome were selected for on LB agar with kanamycin. Colonies were confirmed for vector integration via PCR using insert-specific primers (Table S2). Exconjugants were streaked on LB with kanamycin for purification, then on sucrose-containing LB without NaCl to promote excision of the *sacB*-kanamycin suicide vector and the wild type allele, yielding deletion mutants. Kanamycin-sensitive, sucrose-resistant colonies were PCR-verified for deletion events using flanking confirmation primers (Table S2) and further verified by sequencing.

### Mitomycin C induction, tailocin preparation and normalization

To induce tailocin production in *X. nematophila* strains as well as *P. luminescens* TT01, 100 ml LB was inoculated with 5 ml overnight cultures at OD_600_ of 0.05. Cultures were then grown to an OD_600_ of 0.5-0.6, at which point tailocin production and release was induced using mitomycin C at a final concentration of 5 µg/ml followed by incubation at 30°C with shaking for 20 h. RNase A and DNase I were added at final concentrations of 1 µg/ml, and cultures were incubated for 30 min at 37°C with shaking, before centrifugation at 3,500 x ***g*** for 20 min to pellet cellular debris. Supernatants were filtered through 0.45 um pores and incubated at 4°C for 4 h at a final concentration of 1 M NaCl and 10% polyethylene glycol (PEG) 8000. Following incubation, filtrates were centrifuged at 13,750 x ***g*** for 15 min. Supernatants were poured off and trace amounts were aspirated with a pipette. Precipitates were resuspended in 3 ml LB and centrifuged again at 6000 x ***g*** for 5 min. The final solution was filtered through 0.2 µm pores and its protein concentration determined using Pierce 660 nm Protein Assay Reagent. Tailocin preparations were cryoprotected with glycerol (4%(v/v)) before storage at 4°C.

### Tailocin visualization by transmission electron microscopy

For transmission electron microscopy (TEM) preparation, 200 µl of tailocin preparation was normalized to an A_280_ reading of 0.15, then ultracentrifuged at 287,500 x ***g*** for 15 min. The supernatant was removed, and pellets were resuspended in 50 mM Tris-HCl, pH 8.7. A TEM negative staining protocol was followed. Briefly, a carbon coated copper grid (CFT200 -Cu-50, Electron Microscopy Science) was placed over a 20 µl drop of tailocin sample for a minute. Excess sample was dabbed off the grid using filter paper, prior to rinsing by dipping the grid in distilled water before removing excess distilled water using filter paper. The grid was then stained with Uranyless (Electron Microscopy Sciences) for a minute and air dried. Imaging was performed using a JEOL 1400Flash TEM (JEOL USA, Peabody, Massachusetts), and all images captured at 120kV. Individual tailocin particles were visualized and their images processed using ImageJ software. The lengths of the extended tailocins were taken from the baseplate to the collar and scale bars of 100 nm in length were added for measuring purposes.

### Growth inhibition assays of mitomycin-induced cell lysates from *X. nematophila*

To assay growth inhibition, overnight cultures were sub-cultured to an OD_600_ of 0.05 in 5 ml LB then grown to an OD_600_ of 0.5-0.6 and diluted 1:750. In a 96 well plate, 100 µl of diluted culture was combined with 50 µl of tailocin preparation (0.25 mg/ml). For negative controls, 50 µl of LB with 4% (v/v) glycerol was used instead of tailocin preparations. All inhibition assays were conducted in triplicate technical replicates. See Supplementary File 2 for biological replicate number for each actor strain. The equation used to calculate percentage growth inhibition is = (1-(OD treatment/OD control)) *100 (Supplementary File 2). For most comparisons, the time point selected for the OD reading was when the control treated culture first plateaued in stationary phase. However, in some cases (e.g., *X. nematophila* ATCC19061 *xnp1* mutant strain tailocins in Fig. 6B) different time points were selected to more accurately represent inhibitory phenotype differences from wild type (Supplementary File 2).

### Growth inhibition assays of mitomycin-induced cell lysates from *X. bovienii*

To determine the killing and susceptibility spectra of 42 *X. bovienii* strains (Supplementary File 1), target strains were assayed for inhibition by mitomycin-induced cell lysates (MILs) from actor strains. Briefly, 5 ml LB broth (Difco) in 20 ml culture tubes was inoculated with individual colonies of each strain picked from freezer stocks. Cultures were grown overnight at 28°C and then used to inoculate LB broth, wherein bacteria were cultured up to an OD_600_ of 0.5, at which point 0.5 µg/ml mitomycin C (Sigma-Aldrich) was added to induce both tailocin and phage production and release through cell lysis (19, 36, 37). After overnight incubation at 28°C, mitomycin-induced lysates (MILs) were obtained by centrifuging cultures at 4,472 x ***g*** for 15 min then filtering the supernatant through 0.45 µm pore-sized membranes (Acrodisc) and filtrates were stored at 4°C until their use in testing.

The inhibitory activity of each MIL was tested by spotting 10 μl onto soft nutrient agar (5 g/L) sowed with 2% (v/v) of stationary-phase liquid culture of a recipient colony. Plates were incubated for 48 h at 28°C, at which timepoint inhibition could be visualized as a clear zone on the recipient lawn. Assays were scored on a 0-3 scale, with 0 indicating no sign of inhibition, and averaged between two observers. Supernatant from blank LB cultures treated with mitomycin and self-tests, whereby both the MIL and the recipient colony were of the same strain, were used as negative controls. Two independent inductions were performed for each strain and each actor was tested against at least 16 recipients. Recipient strains also included unsequenced strains of *X. bovienii* and *X. koppenhoeferi*. Replicate assays for the same actor-recipient pair were averaged. Using SAS v. 9.4, the calculation of phenotypic distances between killing profiles for each strain was performed using the Euclidean method, while hierarchical clustering was performed using Ward’s Minimum Variance Method. Resistance profiles were calculated and clustered similarly, based on recipients that were tested on at least two different assay dates and against a minimum of 12 actors. In total, 32 sequenced strains were characterized for their killing, and 34 strains for their susceptibility based on 1,915 inhibition tests.

## ACKNOWLEDGEMENTS

Work in the HGB lab was supported by the University of Tennessee David and Sandra White Endowed Fund, the Tennessee Center of Excellence Science Alliance (SA) Joint Directed Research Development (JDRD) program, and the US National Science Foundation EDGE grant (IOS-2128266). ECA was supported by funds from the University of Tennessee Oak Ridge Innovation Institute and AT was supported by funds from the National Science Foundation Site Award (DBI-2050743). Work in the FB lab was supported by Indiana University and NSF grants DEB 0919015 and 0515832. The authors deeply appreciate the work of Jaydeep Kolape and the UTK Advanced Microscopy Imaging Core for their technical support in transmission electron microscopy. The authors especially wish to thank H. Hawlena and C. Lively for their indispensable effort in collecting and characterizing the *X. bovienii* strains. Additional logistical support was provided by A. Horwitz, C. Collins, C. Syverson, E. Betz, S. Young and R. Mateson. The authors acknowledge and thank Dr. Daren Ginete and Elizabeth Ransone for their early work on the relationship between the phage-related mobilome and strain diversification among *Xenorhabdus bovienii* strains, and Dr. Steven Forst for his work on the elucidation of the *xnp1* locus and for sharing *X. nematophila* AN6/1. The authors thank S. Patricia Stock for sharing *S. anatoliense* and *S. websteri* nematodes from which *X. nematophila* Anatoliense and Websteri were isolated, Jae-Ho Shin for sharing *X. nematophila* C2-3, and Patrick Tailliez for sharing *X. nematophila* F1.

## SUPPLEMENTARY FIGURES AND TABLES

**Figure S1.**
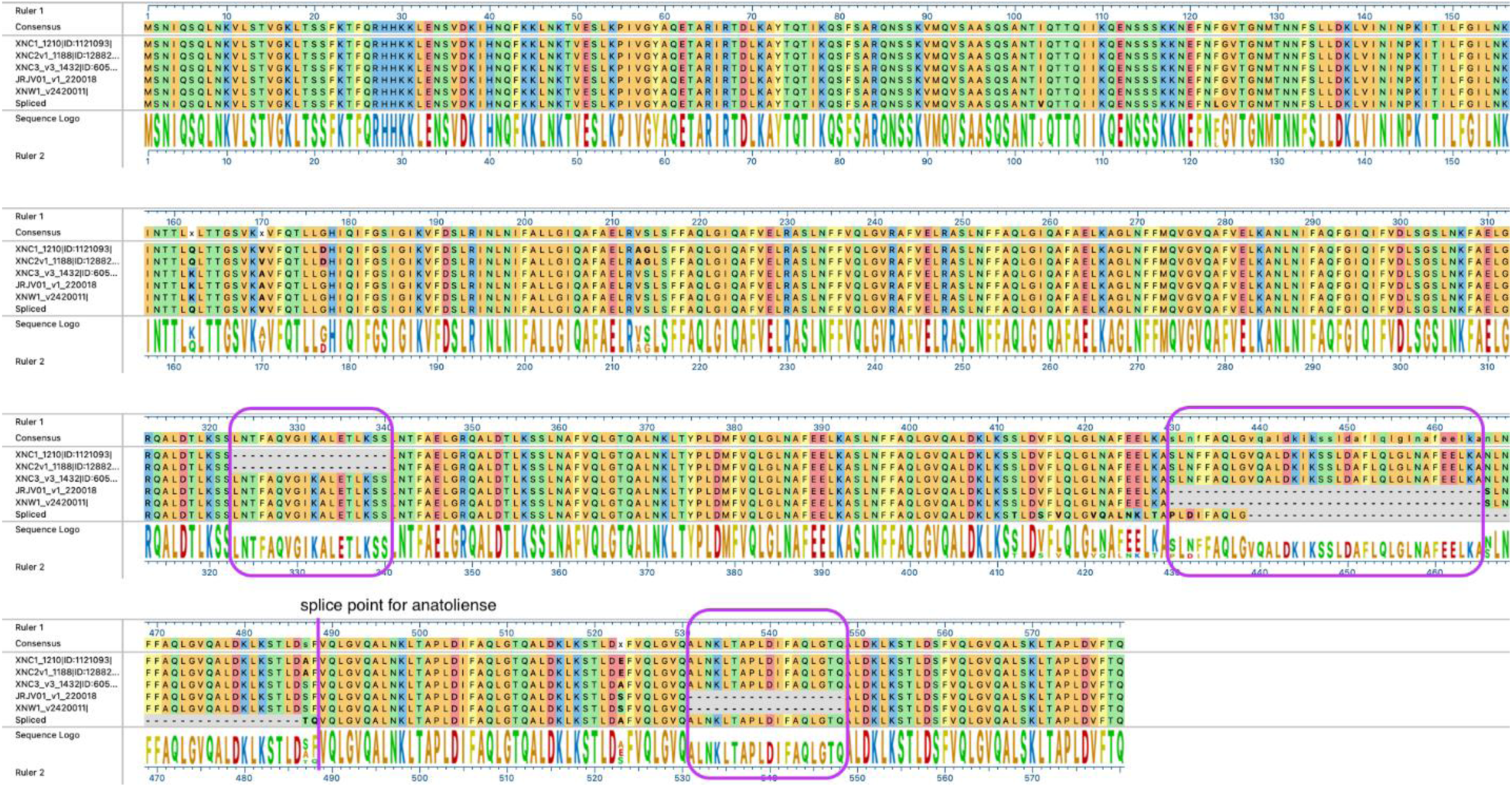
*X. nematophila* predicted *xnp1* locus-encoded tape measure proteins do not vary dramatically in length. Purple boxes indicate regions in which one or more tapemeasure proteins lack amino acids conserved in the others. The predicted tapemeasure coding region for *X. nematophila* Anatoliense spanned a contig break, so the two open reading frames were spliced together for comparison purposes. The six sequences were similar across the full length of the protein with amino acid identities ranging from 98.13% to 100%.

**Table S1.** Strains and plasmids used in this study.

| Xenorhabdus nematophila strains |  |  |  |  |  |  |
| --- | --- | --- | --- | --- | --- | --- |
| Strain | Genotype | HGB number | Nematode host | Geographic origin | Genome Accession/Locus tag | Source/Reference |
| ATCC19061 | Wild type | HGB800 | S. carpocapsae<br>Weiser DD-136/All | USA | GCF_000252955.1-RS_2024_03_28 | ATCC |
| | $\Delta xnpH1$ | HGB2724 | | | XNC1_1193 | This study |
| | $\Delta xnpS1$ | HGB2766 | | | XNC1_1206 | This study |
| | $\Delta xnpH1$<br>$\Delta xnpS1$ | HGB2767 | | | | This study |
| Anatoliense | Wild type | HGB1418 | S. anatoliense | Turkey | GCF_000820905.1 | P. Stock; Hazir *et al.*, 2003 |
| | $\Delta xnpH1$ | HGB2762 | | | XNA1_v220011 | This study |
| AN6/1 rif<sup>R</sup> | Wild type | HGB081 | S. carpocapsae | USA | GCF_000953355.1 | S. Forst |
| F1 | Wild type | HGB1849 | S. carpocapsae<br>Plougastel | France | GCF_000389595.1 | P. Tailliez; Lanois *et al.*, 2013 |
| C2-3 | Wild type | HGB2299 | S. carpocapsae | Republic of Korea | GCF_000785665.1 | S.-J. Hun and J.-H. Shin; Hong *et al.*, 2015 |
| Websteri | Wild type | HGB1419 | S. websteri | Peru | GCF_000820925.2 | S.P. Stock; Nguyen 2007 |
| Other bacterial strains |  |  |  |  |  |  |
| Lab Strain Number |  | Description/Use |  |  |  | Source/Reference |
| HGB1261 | | BW29427 *Escherichia coli* S-17 $\lambda$ pir dap-dependent strain | | | | K.A. Datsenko and B.L. Wanner |
| HGB0801 |  | *Photorhabdus luminescens* TT01 |  |  |  | András Fodor |
| Plasmids |  |  |  |  |  |  |
| Name |  | Description/Use |  |  |  | Source/Reference |
| pKanWor |  | pBluescript II KS+ (Stratagene) with Km cassette (1 Kb) in BamHI site. HGB0181 |  |  |  | E. Vivas and H. Goodrich-Blair |
| pKNGKan |  | pKNG101 derivative with the Str cassette replaced by a Km cassette from pKanWOR. HGB1373 |  |  |  | E. Herbert and H. Goodrich-Blair |
| pKNGKan -ΔXNC1_xnpH1 | | pKNGKan carrying *X. nematophila* ATCC19061 $\Delta xnpH1$ gene construct flanking the multiple cloning site. HGB2723 | | | | This study |
| pKNGKan -ΔXNC1_xnpS1 | | pKNGKan carrying *X. nematophila* ATCC19061 $\Delta xnpS1$ gene construct flanking the multiple cloning site. HGB2765 | | | | This study |
| pKNGKkan -ΔXNA1_xnpH1 | | pKNGKan carrying *X. nematophila* Anatoliense $\Delta xnpH1$ gene construct flanking the multiple cloning site. HGB2761 | | | | This study |

**Table S2.** Xenorhabdicin tapemeasure proteins:

| Strain | *xnp1* tapemeasure locus tag | Predicted length of *xnp1*-<br>encoded tapemeasure<br>protein (aa) |
| --- | --- | --- |
| 19061 | XNC1_1210 | 1036 |
| AN6/1 | XNC2v1_1188 | 1036 |
| F1 | XNC3_v3_1432 | 1054 |
| C2-3 | JRJV01_v1_220018 | 1000 |
| Anatoliense | XNA1_v21110010 | 1006 (spliced) |
|  | XNA1_v21720001 |  |
| Websteri | XNW1_v2420011 | 1000 |

**Table S3.**
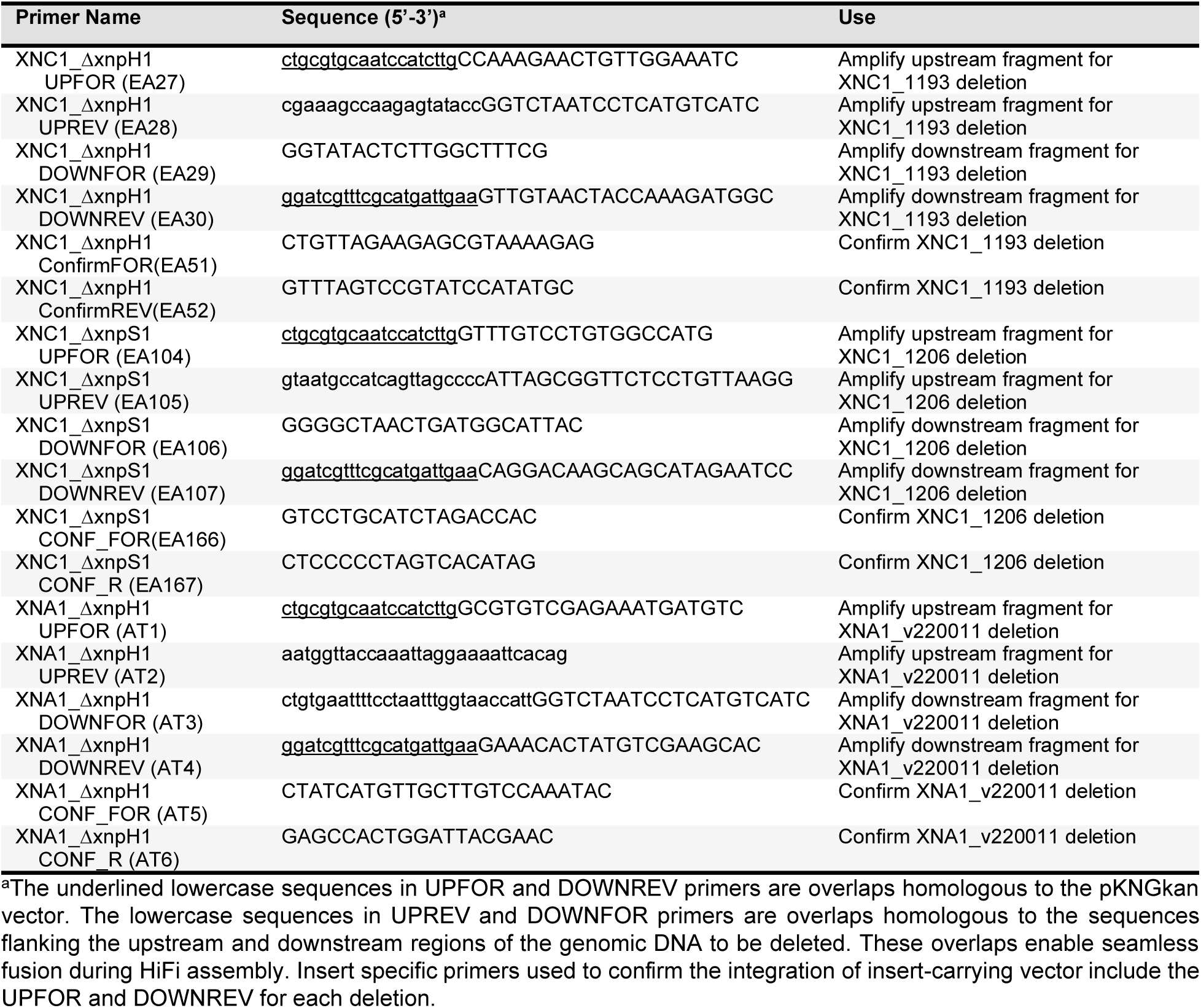
Primers used in this study.

## SUPPLEMENTARY FILES

Supplementary File S1. Genomic analyses and *Xenorhabdus bovienii* strain information

Supplementary File S2. *Xenorhabdus nematophila* tailocin killing assay data

